# A novel tolerance index to identify heat tolerance in cultivated and wild barley genotypes

**DOI:** 10.1101/2020.05.31.125971

**Authors:** Forouzan Bahrami, Ahmad Arzani, Mehdi Rahimmalek

## Abstract

Thermal stress at the reproductive stage poses a substantial constraint on cereal production worldwide. This study was conducted to assess tolerance to terminal high-temperature stress in 45 wild (*Hordeum vulgare* ssp. *spontaneum)* genotypes, 4 cultivars (*H. vulgare* ssp. *vulgare*), 98 F3 and 79 BC1F2 families derived from hybridization of the most tolerant wild genotype and a susceptible cultivar ‘Mona’. Results of analysis of variance showed the significant genotypic and high-temperature stress effects on all the traits studied. In contrast to the cultivated genotypes, the wild ones were found less affected by high-temperature stress. The multivariate analysis highlighted the additional high-temperature tolerance components in the tolerant families and wild genotypes. Grain yield strongly correlated (*p* < 0.01) with stress tolerance, yield stability, and heat tolerance indices. The reduction in the reproduction period caused by high-temperature was much higher in cultivated genotypes than in wild ones. In conclusion, the ingenuous-focused strategies like escape/avoidance are being used primarily to cope with heat stress by cultivars, while adaptive-focused coping strategies such as tolerance are being implemented by wild barley.

## 1 | INTRODUCTION

Among the enormous global challenges facing plant breeders is genetic erosion as the one of utmost importance. There is, therefore, an urgent need to exploit genetic resources as an effective solution to enhance adaptation of plants to changing environments and food security. Successful crop improvement for high-temperature tolerance mainly relies on the availability of genetic variability, identification of the genetic and physiological mechanisms involved in high-temperature tolerance, reliable screening approaches, efficient genetic manipulation of desired genetic materials, and ultimate development of heat-tolerant cultivars with suitable agronomic and quality attributes (Arzani & Ashraf, 2016). As an immediate progenitor of barley (*Hordeum vulgare* ssp. *vulgare* L.) cultivars, wild barley, the *H. vulgare* ssp. *spontaneum* L. (hereafter referred to as *H. spontaneum*) species is believed to be drought and heat-tolerant (Hubner et al., 2009; Bahrami et al., 2019; Arzani & Ashraf, 2016). Hence, introgression of the wild genome into domesticated barley is therefore of interest for the genetic improvement of abiotic stress tolerance in cultivated barley and to investigate the tolerance mechanisms.

The global warming phenomenon leads to an increase in climatic instability that adversely affects ecosystem quality, plant growth, and agricultural production (Schauberger et al., 2017; Hassan et al., 2020). Terminal high-temperature stress is a term referring to the progressive rise of temperature during the day and in consecutive days during the reproduction stage in both spring and winter type cereals (Bahrami et al., 2019). Grain yield reductions due to high temperatures during the grain-filling stage were found in such cereals as wheat (Dwivedi et al., 2017), barley, and oat (Klink et al., 2014). Terminal heat-tolerant genotypes of *H. spontaneum* have already been found to employ physiological strategies to maintain grain yield under high-temperature stress, such as retention of chlorophyll pigment and protection of photosynthetic apparatus (Bahrami et al., 2019).

Effective selection of plants tolerant to abiotic stresses (drought, salinity, heat, and cold) depends upon the plant breeder’s ability to find a reasonable compromise among grain yield under stress conditions, yield losses due to stress, and yield stability. Therefore, there is a growing need for effective selection indices to explore the phenotypes that correspond to possible genotypes in environmental plasticity and adaptation. The index must ensure plant breeders they can safely select genotypes not only of high yield in optimal conditions but also with reasonably consistent yields for cultivation under stressful conditions. This is while, drought tolerance indices are used to assess high-temperature tolerance in crop plants such as wheat (Hassan et al., 2016; Aziz et al., 2018), durum wheat (Kamrani et al.,2017), and common bean (Chavez-Arias et al., 2018; Porch, 2006).

In view of the above-mentioned considerations, the current study was designed and conducted: 1) to evaluate the agronomic responses of *H. spontaneum* germplasm originated from west of Iran to high-temperature stress during the grain-filling period, 2) to determine the most appropriate agronomic traits and yield-based tolerance indices for screening and selecting within BC1F2 and F3 lines for high-temperature tolerance and 3) to describe the relationships among agronomic traits and selection indices of tolerance to high-temperature stress.

## 2 | MATERIALS AND METHODS

### 2.1 | Experimental site, soil moisture, and air temperature

Two independent experiments were conducted at the research farm of Isfahan University of Technology, Lavark, Najaf-Abad, Iran (40 km south-west of Isfahan, 32° 30’ N, 51° 20’ E, 1630 m asl). The field site is characterized by mean annual temperature and precipitation of 14.5°C and 140 mm, respectively, with a silty clay loam soil (typic Haplargids). The air temperatures at reproductive growth period in each of the three years studied, obtained from a nearby meteorological station (3 km) located Najaf-Abad, Iran, are presented in Table 1. The field was irrigated to minimize the confounding effects of drought stress. The irrigation water was applied from a pumping station to the plots through polyethylene pipes equipped with water flow meters. The plots were irrigated when the soil moisture reached above 90% of field capacity (ψ = −0.02) in the root zone. The soil moisture was monitored by taking soil samples at a depth of 0-30 cm. The amount of irrigation water was determined according to the following equation:

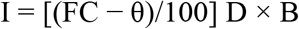

**TABLE 1.** Air temperature of experimental site [mean (± SE)] at reproductive growth stage of barley plants under normal and delayed sowing in 2016, 2017, 2019

| Growing period | Field conditions | Duration/<br>date | Temperatures (°C) |  |  |
| --- | --- | --- | --- | --- | --- |
|  |  |  | Minimum | Maximum | Mean |
| Anthesis | Normal | 4.4.2016-20.4.2016 | 7.5 (0.65)♣ | 22.5 (0.32) | 15.0 |
|  |  | 4.4.2017-19.4.2017 | 6.8 (0.97) | 20.4 (0.72) | 13.6 |
|  |  | 3.4.2019-21.4.2019 | 6.4 (0.39) | 19.5 (0.22) | 13.1 |
|  | Stress | 3.5.2016-13.5.2016 | 17.3 (0.69) | 29.7 (0.16) | 23.5 |
|  |  | 3.5.2017-14.5.2017 | 19.4 (1.07) | 30.1 (0.21) | 24.7 |
|  |  | 1.5.2019-16.5.2019 | 19.6 (0.35) | 29.6 (0.47) | 24.6 |
| Grain-filling | Normal | 21.4.2016-20.5.2016 | 17.4 (1.65) | 25.9 (1.24) | 21.6 |
|  |  | 18.4.2017-20.5.2017 | 18.2 (0.7) | 28.3 (0.63) | 23.2 |
|  |  | 20.4.2019-19.5.2019 | 17.7 (0.83) | 27.8 (1.28) | 22.7 |
|  | Stress | 14.5.2016-8.6.2016 | 22.9 (0.91) | 36.2 (0.47) | 29.5 |
|  |  | 15.5.2017-10.6.2017 | 24.6 (0.77) | 35.6 (0.33) | 30.1 |
|  |  | 13.5.2019-14.6.2019 | 23.8 (0.55) | 35.5 (0.44) | 29.6 |

I is irrigation depth (cm), FC is soil gravimetric moisture percentage at field capacity, θ is soil gravimetric moisture percentage at irrigating time, D is the root-zone depth, and B is the soil bulk density at root-zone (1.4 g/cm3).

### 2.2 | Plant materials and growth conditions

#### 2.2.1 | Experiment 1

Forty-nine barley genotypes comprising 45 *H. spontaneum* genotypes originated from western Iran (i.e., the five provinces of Lorestan, West Azarbaijan, Ilam, Kordestan, and Kermanshah) and the four barley cultivars of ‘Mona’, ‘Reyhan’, ‘Nosrat’, and ‘Fajr30’ were used in this study. A lattice-square design (7 × 7) replicated twice was used in normal and in delayed sowing conditions in 2015-16 and 2016-17 growing seasons. The seeds were sown on two dates, November and January (normal and delayed, respectively), in two consecutive years (2015and 2016). Each plot consisted of three rows, 4 m long and 30 cm apart. At maturity, barley plants from 3 m of the middle row in each plot were harvested to determine grain yield and its components.

#### 2.2.2 | Experiment 2

The interspecific hybrids (F1) were produced from a cross between a high-temperature tolerant genotype of *H. spontaneum* and a barley cultivar (*H. vulgure* var. ‘Mona’). The crossing protocol is summarized as follows: 1) all 45 wild genotypes were crossed with the ‘Mona’ cultivar in 2015-16, 2) the F1 plants were both selfed and backcrossed to generate F2 and BC1F1, using the ‘Mona’ cultivar as the recurrent parent, 3) the two-year data (2015-17 growing seasons) from experiment 1 were used to identify the most high-temperature tolerant wild genotype (genotype no. 2); subsequently, the F2 and BC1F1 plants derived from the cross between this wild genotype and ‘Mona’ were subjected to selfing, and the resultant 98 F3 and 79 BC1F2 families were used in the experiment 2. These lines were grown along with their parents (wild barley no.2 and ‘Mona’) in two planting dates (normal: November, and delayed: January) of 2018-19 growing season under field conditions. Two separate 10 × 10 and 9 × 9 lattice-square designs with two replications were used for F3 and BC1F2 families, respectively. Each line was sown in a two-row plot of 2 m length with a row spacing of 30 cm. At maturity, plants in 1 m of the middle of the two rows in each plot were harvested to determine grain yield and its components.

### 2.3 | Measurement of traits

#### 2.3.1 | Agronomic traits

Days to anthesis (DA) was defined as the number of days from the planting date until one-half of the plants had at least one extruded anther. Days to physiological maturity, hereafter referred to as ‘days to maturity’ (DM), was measured as the number of days from the planting date to the appearance of peduncle senescence in the whole plot. Moreover, the length of the reproductive growth period (RGP) was calculated by DA subtracted from DM. Spike length (SPL), plant height (PH), number of fertile tillers (NT), and number of grain per spike (NGS) were recorded at the physiological maturity stage on 10 plants per plot. The average weight of 2 samples of 1000 grains randomly taken from each plot was used to determine 1000 grain weight.

#### 2.3.2 | Physiological traits

The following physiological traits were measured in F3 and BC1F2 families:

Chlorophyll content (Chla and Chlb) and carotenoid content (Car) were extracted based on the standard method of Lichtenthaler and Wellburn (1983) using acetone 80% from fresh leaves sampled at the grain-filling stage. Concentrations were measured according to the absorbance of the solutions at 663, 646, and 470 nm and expressed as mg per g fresh leaf sample (mg g^-1^ FW) using the following equations:

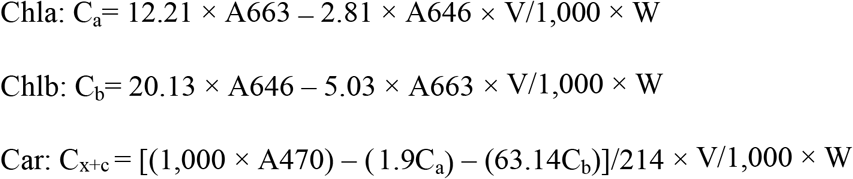

Where V is the volume of acetone (ml), and W is the fresh weight (g).

Then total chlorophyll content (Chla+b) and the ratio of Chla to Chlb (Chla/b) were calculated.

Proline content (Pro) was determined using fresh leaves sampled at the grain-filling stage. Fresh leaf (0.2 g) powdered in liquid nitrogen was homogenized in 10 ml of 3% aqueous sulfosalicylic acid then centrifuged for 10 min at 10,000 rpm. Subsequently, proline content was calculated according to Bates et al. (1973) method and expressed in mg g^-1^ of fresh weight (FW).

#### 2.3.3 | Tolerance indices

Tolerance indices including tolerance index (TOL) (Rosielle & Hamblin, 1981), stress tolerance index (STI) (Fernandez, 1992), stress susceptibility index (SSI) (Fischer & Maurer, 1978), yield stability index (YSI) (Bouslama & Schapaugh, 1984) were calculated using the equations given below.

1. Tolerance index (TOL)

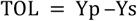
2. Stress tolerance index (STI)

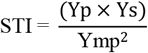
3. Stress susceptibility index (SSI)

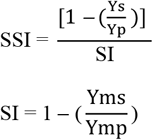
4. Yield stability index (YSI)

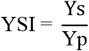 Yp and Ys represent the mean yields of each genotype in normal and high-temperature stress conditions, respectively. While, Ymp and Yms denote the mean yields of all genotypes in normal and high-temperature stress conditions, respectively. In this paper, a modified version of the tolerance index named hereafter as heat tolerance index (HTI), was proposed. In this equation, HTI for each genotype is calculated by dividing Ymp and Yms to Yp and Ys, respectively, and then subtracting the resulting products as below:
5. Heat tolerance index (HTI)

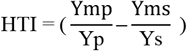

### 2.4 | Statistical analysis

A combined analysis of variance (ANOVA) was carried out for the traits studied using the GLM procedure of SAS software (Version 9.3; SAS Institute 2011). Also, the efficiency of the lattice design was compared (tested) with that of the randomized complete block design (RCBD) based on the ratio of lattice mean square of residuals (error) × 100/RCBD mean square residual for each trait. Due to the slight differences between the two, the RCBD design was used for data analysis in both experiments. Comparison of means was performed using the Fisher’s least significant difference (LSD5%) test. Multivariate statistical analysis comprising calculation of phenotypic correlation (Pearson coefficients) and principal component analysis (PCA) employed for the traits studied and the tolerance indices for both experiments (*n* = 49 genotypes (45 wild + 4 cultivars), *n* = 98 F3 families, and *n* = 79 BC1F2 families). Biplot graphs based on PCA were prepared using Stat Graphics. The Sigma Plot software was then used to develop a three-dimensional graph for identifying superior genotypes and families based on the best tolerance indices identified.

## 3 | RESULTS

As shown by the data summarized in Table 1, average daily minimum and maximum temperatures have increased during the grain-filling period in delayed sowing when compared to the normal sowing. In normal and high-temperature stress conditions, the overall mean of maximum temperature during grain filling in the three growing seasons (2015-2019) was increased from 27.3 to 35.8, while the mean of minimum temperature changed from 17.8 to 23.8 °C, respectively.

### 3.1 | Evaluation of wild and cultivated barley genotypes

The analysis of variance showed that terminal high-temperature had significant effects on all the traits studied (Table 2). Moreover, highly significant (*p*<0.01) variations in phenological and agronomic traits were found in wild and cultivated genotypes. The two growing years (Y) showed significant differences only for the NGS and GW traits. The interaction between genotype and environment (G×E) was also significant for all the traits except DA, SPL, and NGS. Furthermore, the environment by year (E×Y) interaction was significant for PH and yield components.

**TABLE 2.**
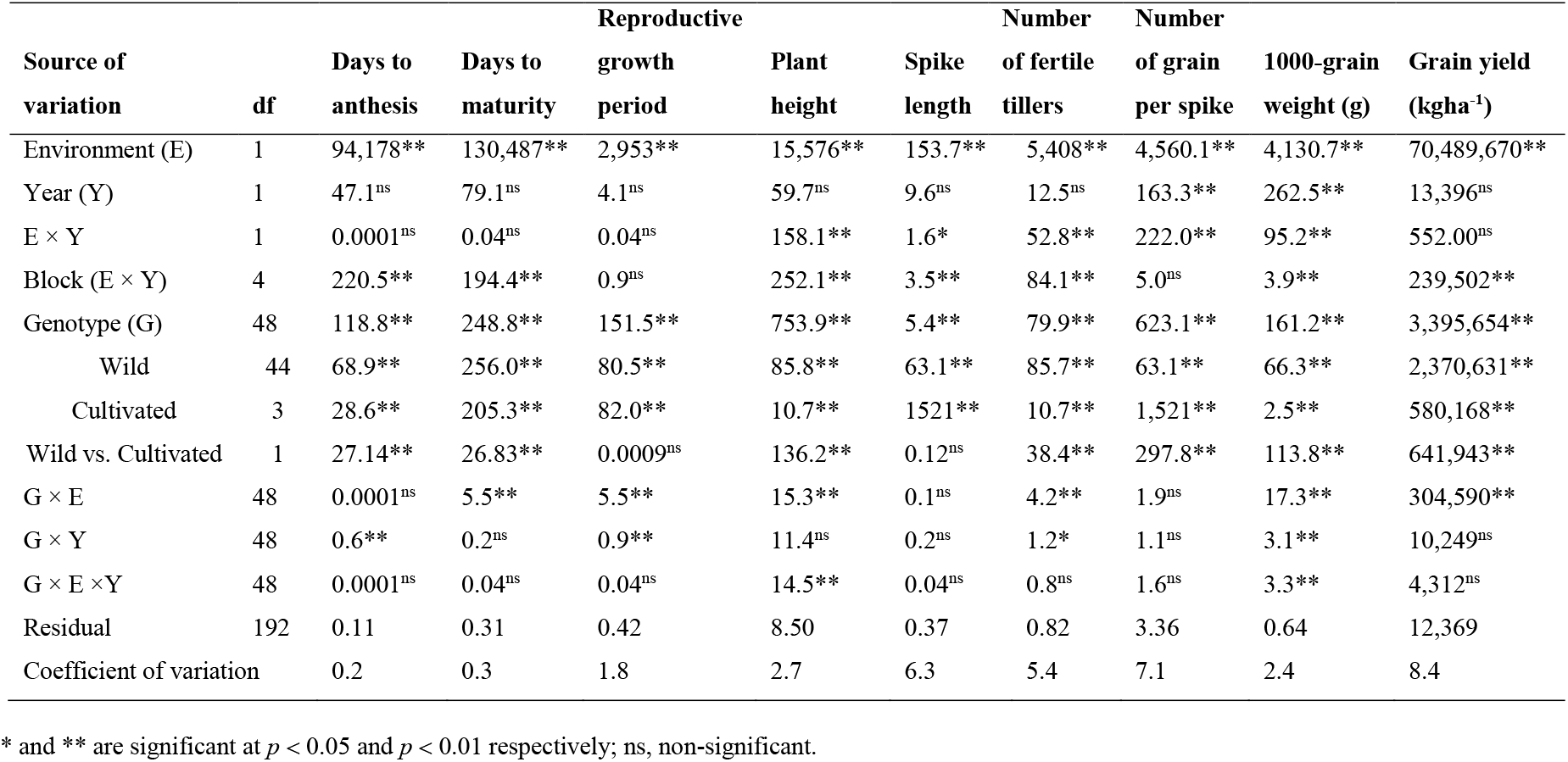
Mean squares of combined analysis of variance for phenological and agronomic traits in wild and cultivated barley genotypes grown under normal and high-temperature stress conditions in two consecutive years (2016 and 2017)

Mean comparisons of the agronomic traits averaged over the harvested years of 2016 and 2017 in high-temperature stress conditions indicated a high genetic variation among the wild genotypes. The wild genotype no. 2 produced the highest yield and yield components among 45 wild genotypes. The lines derived from hybridization between this wild genotype and ‘Mona’ have been used in our experiment 2 to maximize accuracy power for identifying agronomic attributes and indices relevant to sustainable production under normal and high-temperature stress conditions. In comparison to normal conditions, high-temperature stress caused a significant reduction in all the traits evaluated, including grain yield, in both wild and cultivated genotypes (Table 3). Moreover, significant differences were observed between wild and cultivated genotypes for all the traits studied except for DM and SPL under both environments. DA was longer for the wild genotypes than the cultivars, while longer RGP was found for the cultivars. Despite the longer RGP in the cultivated barley, they exhibited a two-fold reduction in RGP as a result of high-temperature stress compared with wild ones (Table 3). In contrast to the cultivated genotypes, the wild ones were less affected by terminal high-temperature in their NGS and GW (Table 3). Barley cultivars differed significantly from wild genotypes in terms of yield and its components in both environmental conditions (Table 3).

**TABLE 3.** Means (±SE) of phenological and agronomic traits separately in wild and cultivated barley genotypes grown under normal and high-temperature stress conditions in two consecutive years (2016 and 2017)

| Traits | Normal |  | Stress |  | Change (%)<sup>††</sup> |  |
| --- | --- | --- | --- | --- | --- | --- |
|  | Wild | Cultivated | Wild | Cultivated | Wild | Cultivated |
| Days to anthesis | 157.13(0.24)<sup>a†</sup> | 147.40(0.59)<sup>b</sup> | 126.13(0.25)<sup>a</sup> | 117.25(0.56)<sup>b</sup> | -19.72 | -20.45 |
| Days to maturity | 194.70 (0.43)<sup>a</sup> | 199.50(1.17)<sup>a</sup> | 159.00(0.43)<sup>a</sup> | 156.75(1.20)<sup>a</sup> | -18.33 | -21.42 |
| Reproductive growth period | 37.57(0.23)<sup>b</sup> | 52.10(0.78)<sup>a</sup> | 32.86(0.24)<sup>b</sup> | 39.50(0.69)<sup>a</sup> | -12.53 | -24.18 |
| Plant height | 114.68(0.47)<sup>a</sup> | 86.18(1.45)<sup>b</sup> | 101.87(0.57)<sup>a</sup> | 75.81(1.77)<sup>b</sup> | -11.17 | -12.03 |
| Spike length | 10.17(0.06)<sup>a</sup> | 10.87(0.39)<sup>a</sup> | 8.84(0.07)<sup>a</sup> | 8.99(0.37)<sup>a</sup> | -13.00 | -17.30 |
| Number of fertile tillers | 19.25(0.25)<sup>b</sup> | 21.93(0.41)<sup>a</sup> | 12.85(0.27)<sup>b</sup> | 13.38(0.53)<sup>a</sup> | -33.24 | -39.00 |
| Number of grains per spike | 26.01(0.22)<sup>b</sup> | 56.50(3.10)<sup>a</sup> | 21.15(0.25)<sup>b</sup> | 40.65(3.23)<sup>a</sup> | -18.18 | -28.01 |
| 1000-grain weight (g) | 34.09(0.23)<sup>b</sup> | 50.16(0.25)<sup>a</sup> | 28.99(0.29)<sup>b</sup> | 36.11(0.40)<sup>a</sup> | -14.95 | -28.11 |
| Grain yield (kg ha<sup>-1</sup>) | 1,417.2(24.4)<sup>b</sup> | 4,247.4(168.1)<sup>a</sup> | 819.0(18.8)<sup>b</sup> | 2,069.1(103.6)<sup>a</sup> | -42.20 | -51.28 |

<sup>†</sup>Mean values of each traits in each environment (normal or high-temperature stress) are not significantly different ( $p < 0.05$ according to LSD test) between wild and cultivated barley, if followed by the same letter .
<sup>††</sup>Minus sign represents decreasing percentage due to high-temperature stress conditions.

Five yield-based tolerance indices were employed to screen superior tolerant wild genotypes for high-temperature stress. Barley cultivars had the highest reduction in grain yield (TOL values) due to high-temperature stress compared to the wild genotypes. Moreover, the wild genotypes were superior for the stability-oriented indices (YSI and HTI). The STI, YSI, and HTI correlated with Ys in the barley genotypes (0.97**, 0.82**, and 0.87**, respectively). In contrast, grain yield had a high association with STI and TOL in the cultivated genotypes (0.99** and 0.90**, respectively) under high-temperature stress.

PCA was carried out using phenological and agronomic traits and tolerance indices for the 49 barley genotypes grown under high-temperature stress conditions. Results showed that the first two components explained 72.08% of the cumulative variance in the traits studied and tolerance indices (Figure 1). The angles between the vectors show the relationships between the variables. TOL, NGS, Yp, STI, and GW had positive correlations, but the correlation between DA and PH was negative under high-temperature stress (Figure 1). Moreover, positive correlations were observed among YSI, SPL, HTI, NT, and DM. The PC1 was termed ‘yield potential’ to reflect the high and positive correlations with grain yield and its components. Accordingly, the high-temperature tolerant wild genotypes are located in the high PC1 and PC2 regions of the biplot, while the heat-sensitive ones are placed in the low PC1 and PC2 regions. Then, accordingly, the genotypes subjected to high-temperature stress conditions were classified into the three groups located in the regions enclosed by circular lines in Figure 1. The first group consisted of genotypes with high PC1 and PC2 values and included those located closest to YSI and HTI; they mostly originated from Illam and Kermanshah provinces. The second group consisted of the three ‘Reyhan’, ‘Nosrat’, and ‘Fajr30’ cultivated barley genotypes. Finally, the third heat-sensitive group (i.e., with the lowest PC1 and PC2 values) consisted of genotypes that had been collected from Kamyaran and Sanandaj regions in Kordestan Province.

**FIGURE 1.**
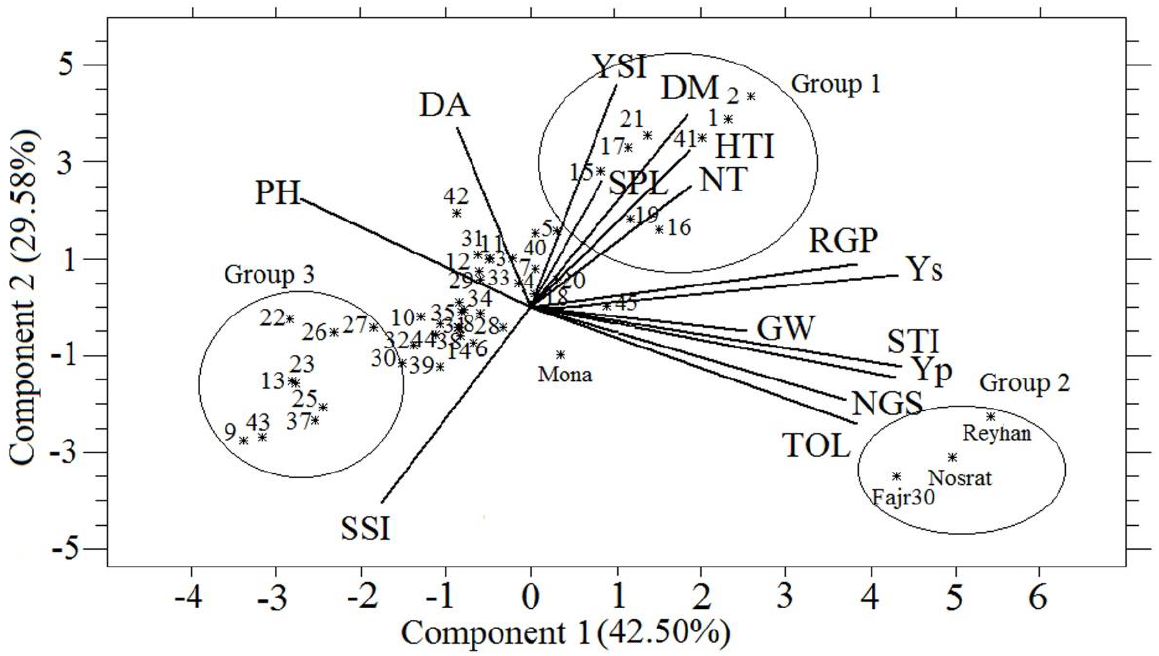
Biplot of PC1 versus PC2 based on the mean values obtained from each trait during the two-year study period for the high-temperature treatment. DA, days to anthesis; DM, days to maturity; GW, 1000-grain weight; HTI: heat tolerance index; NGS, number of grain per spike;NT, number of fertile tillers; PH, plant height; RGP, reproductive growth period; SPL, spike length; SSI, stress susceptibility index; STI, stress tolerance index; TOL: tolerance index; Yp, grain yield under normal conditions; Ys, grain yield under high-temperature stress conditions; YSI, yield stability index. Numbers in the figure are wild genotypes.

Regarding the strong inter-correlations among STI, YSI, and HTI, a threedimensional graph was plotted to illustrate the relationships among the studied wild genotypes based on these variables (Figure 2). The genotypes were placed into three zones (A, B and, D), representing the three tolerance indices. That is, in other words, about one-fourth of the wild genotypes with the highest yield stability and adaptability values located in the ‘A’ zone. Whereas, genotypes in the ‘D’ zone exhibited the highest susceptibility to terminal high-temperature, mostly originated from Lorestan, Kordestan, and West Azarbaijan provinces.

**FIGURE 2.**
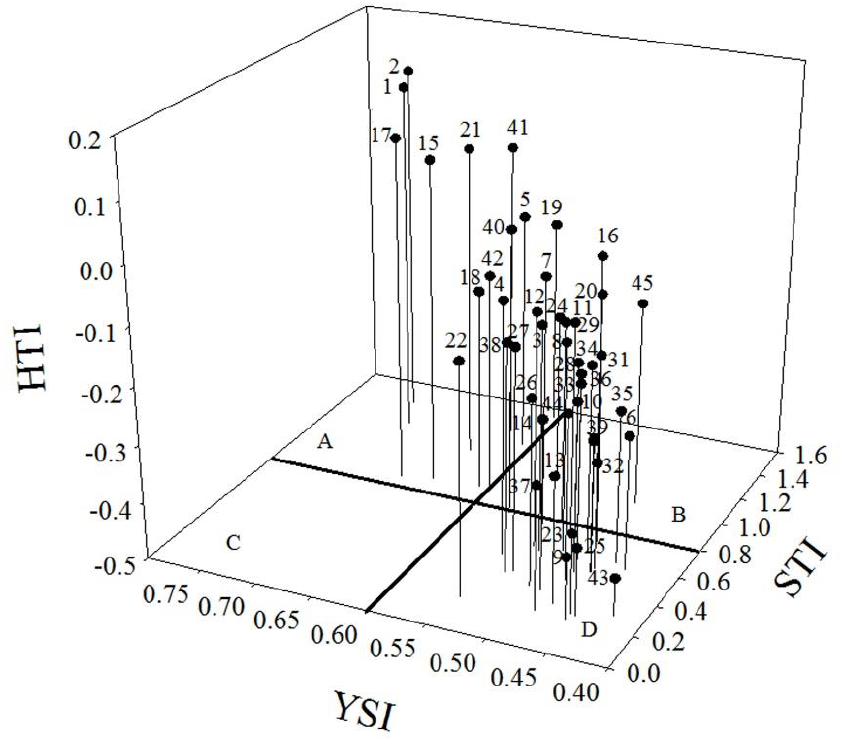
Three-dimensional diagram based on values obtained stress tolerance index (STI), yield stability index (YSI) and heat tolerance index (HTI). Numbers represent wild genotypes.

Multivariate analysis classified the wild genotypes no. 1, 2, 5, 15, 17, 18, 19, 21, 40, and 41as the terminal high-temperature tolerant group.

### 3.2 | Evaluation of F3 and BC1F2 barley families

Analysis of variance showed the significant effect of high-temperature stress on all the traits in both generations except for Chla/b in the F3 families (data not shown). Moreover, the families groups showed a highly significant (*p*<0.01) variation in phonological, agronomic, and physiological traits. In the F3 families, the interaction between genotype by environment (G×E) was not significant for all the traits except for GW, Chla, Chlb, and Chla/b. However, the G×E effect was significant for yield, its components, and Chla/b in BC1F2 families (data not shown).

The response of the families and their parents to high-temperature stress was evaluated using the overall means of the traits (Table 4 and 5). The wild genotypes were a few days later in DM and several days later in DA when compared with the cultivars (Table 3). In contrast, the cultivars had a longer RGP compared with the wild genotypes. In comparison, the progeny lines showed an intermediate maturity time relative to their wild and cultivated parents (Table 4). Under normal and high-temperature stress conditions, the cultivated parent showed the highest SPL, whereas the highest reduction percentage of SPL due to high-temperature stress was recorded for F3 families (Table 4). The lowest reduction for NT was observed in the F3 and BC1F2 generations compared with their parents (Table 4). The lowest and highest reduction for NGS was 24.5% and 10.63% in the F3 generation and cultivated parent. The highest GW and grain yield was observed in cultivated parent under normal conditions. There was no significant difference between BC1F2 families and cultivated parent for GW and grain yield under high-temperature stress. Nevertheless, the wild parent showed the lowest grain yield reduction (Table 4). Moreover, some F3 and BC1F2 families out-yielded their parents, which indicating transgressive segregation for grain yield in these families (data not shown).

**TABLE 4.**
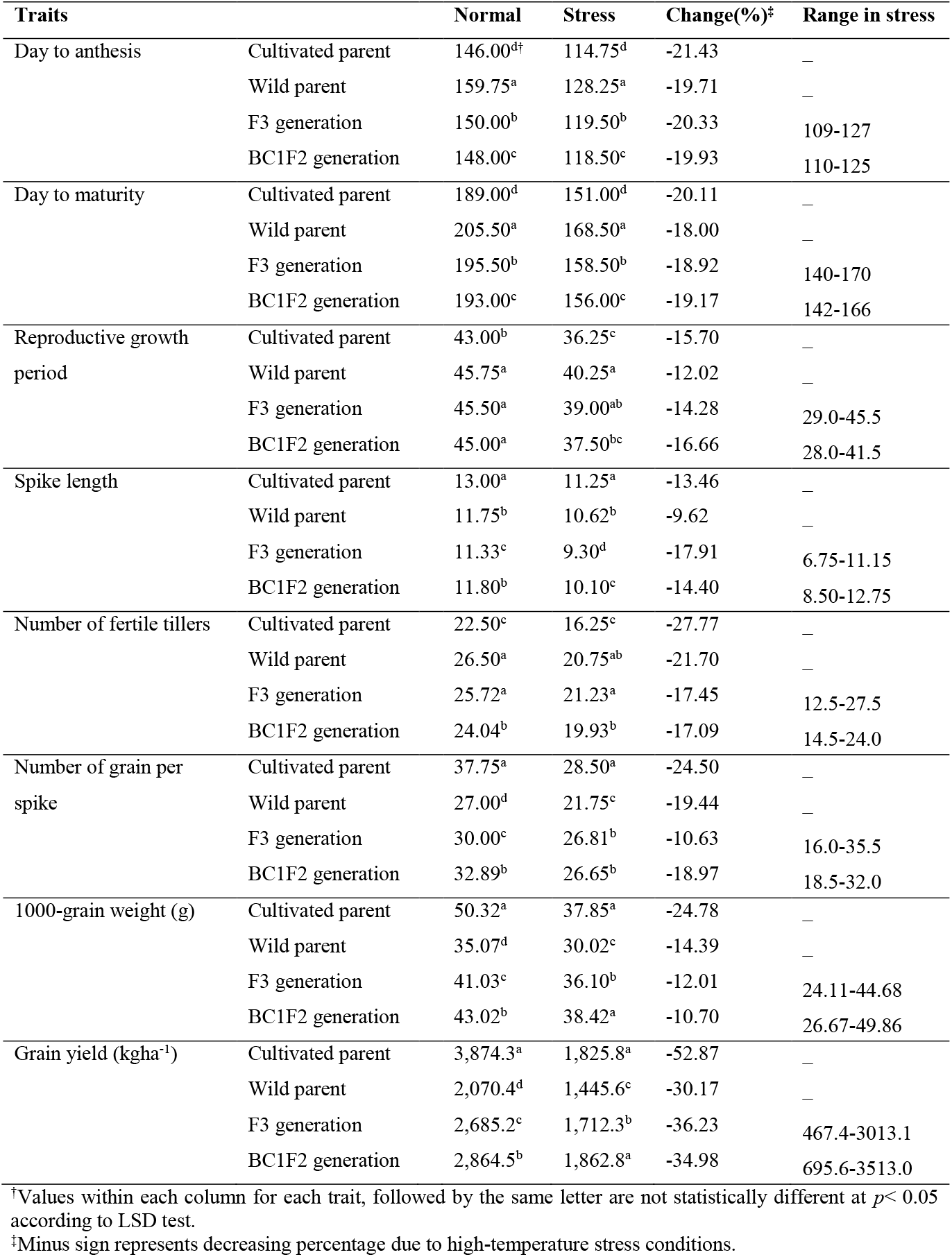
Means of phenological and agronomic traits separately in F3 and BC1F2 generations and their parents grown under normal and high-temperature stress field conditions in 2018-2019

**TABLE 5.** Means of physiological traits separately in F3 and BC1F2 generations and their parents grown under normal and high-temperature stress field conditions in 2018-2019

| Traits |  | Normal | Stress | Change (%)<sup>‡</sup> | Range in stress |
| --- | --- | --- | --- | --- | --- |
| Chlorophyll a<br>(mg g<sup>-1</sup>FW) | Cultivated parent | 1.95<sup>a</sup> | 1.25<sup>a</sup> | -35.89 | – |
|  | Wild parent | 1.61<sup>c</sup> | 1.18<sup>b</sup> | -26.71 | – |
|  | F3 generation | 1.55<sup>d</sup> | 1.05<sup>c</sup> | -32.25 | 0.47-1.25 |
|  | BC1F2 generation | 1.68<sup>b</sup> | 1.16<sup>b</sup> | -31.54 | 0.57-1.35 |
| Chlorophyll b<br>(mg g<sup>-1</sup>FW) | Cultivated parent | 0.97<sup>a</sup> | 0.49<sup>b</sup> | -49.50 | – |
|  | Wild parent | 0.78<sup>b</sup> | 0.45<sup>b</sup> | -42.30 | – |
|  | F3 generation | 0.99<sup>a</sup> | 0.55<sup>a</sup> | -44.44 | 0.11-1.38 |
|  | BC1F2 generation | 0.94<sup>a</sup> | 0.54<sup>a</sup> | -43.61 | 0.13-1.23 |
| Total<br>chlorophyll | Cultivated parent | 2.92<sup>a</sup> | 1.75<sup>a</sup> | -40.06 | – |
|  | Wild parent | 2.39<sup>d</sup> | 1.63<sup>bc</sup> | -31.79 | – |
|  | F3 generation | 2.54<sup>c</sup> | 1.60<sup>c</sup> | -37.00 | 0.94-2.33 |
|  | BC1F2 generation | 2.63<sup>b</sup> | 1.70<sup>ab</sup> | -35.36 | 0.99-2.40 |
| Ratio of<br>chlorophyll a to b | Cultivated parent | 2.01<sup>a</sup> | 2.52<sup>b</sup> | +25.37 | – |
|  | Wild parent | 2.06<sup>a</sup> | 2.62<sup>b</sup> | +27.18 | – |
|  | F3 generation | 1.69<sup>b</sup> | 2.79<sup>ab</sup> | +65.08 | 0.54-12.61 |
|  | BC1F2 generation | 1.95<sup>a</sup> | 3.10<sup>a</sup> | +58.97 | 0.69-10.36 |
| Carotenoid<br>(mg g<sup>-1</sup>FW) | Cultivated parent | 0.75<sup>b</sup> | 0.54<sup>b</sup> | -28.00 | – |
|  | Wild parent | 0.79<sup>a</sup> | 0.59<sup>a</sup> | -25.31 | – |
|  | F3 generation | 0.61<sup>d</sup> | 0.42<sup>c</sup> | -31.14 | 0.24-0.67 |
|  | BC1F2 generation | 0.70<sup>c</sup> | 0.42<sup>c</sup> | -40.00 | 0.27-0.56 |
| Proline<br>(mg g<sup>-1</sup>FW) | Cultivated parent | 0.81<sup>b<sup>c</sup></sup> | 1.85<sup>c</sup> | +128.39 | – |
|  | Wild parent | 1.33<sup>a</sup> | 3.73<sup>a</sup> | +180.45 | – |
|  | F3 generation | 0.74<sup>c</sup> | 2.04<sup>b</sup> | +175.67 | 1.05-3.53 |
|  | BC1F2 generation | 0.87<sup>b</sup> | 2.08<sup>b</sup> | +139.36 | 1.13-3.19 |

<sup>†</sup>Values within each column for each trait, followed by the same letter are not statistically different at $p < 0.05$ according to LSD test.
<sup>‡</sup>Plus and minus signs represent increasing and decreasing percentage due to high-temperature stress conditions, respectively.

Mean comparisons of Chla content showed that cultivated parent had the highest and F3 families had the lowest Chla, under high-temperature stress conditions (Table 5). The cultivated parent had a higher Chla+b and its alteration than the wild parent and two families under both environmental conditions (Table 5). The highest increase for Chla/b due to high-temperature was observed in the F3 and BC1F2 generations (Table 5). Under high-temperature stress, F3 and BC1F2 families had a lower Car value than their parents. Means comparison showed that the proline accumulation due to high-temperature was the highest for wild parent and F3 families (Table 5).

Under high-temperature stress, grain yield was correlated with SPL, NGS, and GW (0.45**, 0.57**, and 0.69**, respectively) in the F3 families. Likewise, a strong positive association was observed between NGS and GW with grain yield (0.56** and 0.72**, respectively) under high-temperature stress in the BC1F2 families. Linear regression analysis showed a highly significant and positive relationship between STI and GW in both generations (Figure 3). The STI, YSI, and HTI correlated strongly with Ys in the F3 families (*r* equal to 0.97**, 0.68**, and 0.58**, respectively). In addition, grain yield showed an association with STI and YSI (0.96** and 0.40**, respectively) under high-temperature stress for BC1F2 generation. A strong association was observed between Pro with grain yield and STI within both generations under terminal high-temperature (Table 6).

**FIGURE 3.**
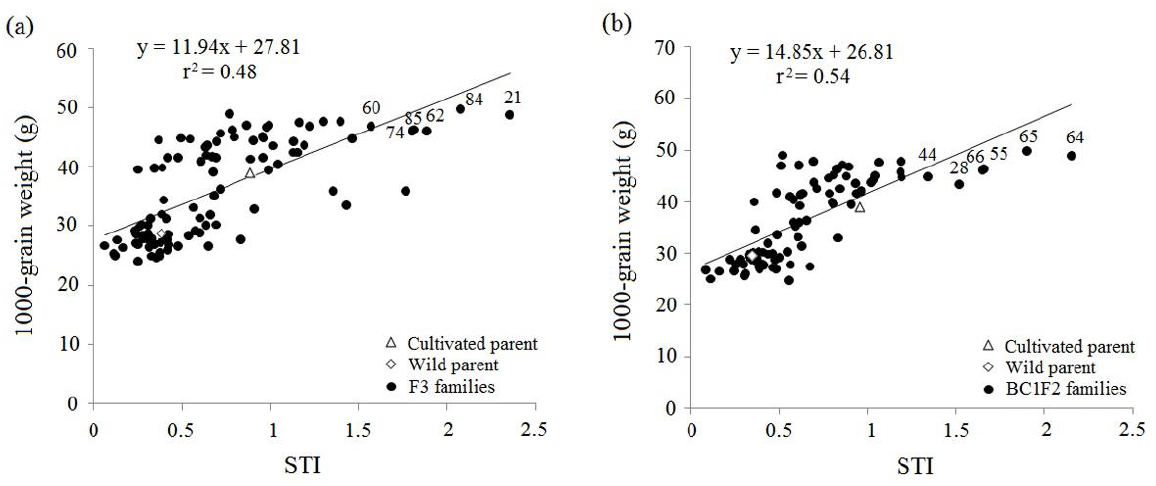
Relationship between 1000-grain weight and stress tolerance index (STI) of F3 (a) and BC1F2 (b) barley families along with their parents under high-temperature stress field conditions.

**TABLE 6.** Correlation coefficients between physiological traits and tolerance indices with in F3 and BC1F2 families grown under high-temperature stress conditions in 2018-2019

| Family | Traits | Chla<br>(mg g<sup>-1</sup><br>FW) | Chlb<br>(mg g<sup>-1</sup><br>FW) | Chla+b | Chla/b | Car<br>(mg g<sup>-1</sup><br>FW) | Pro<br>(mg g<sup>-1</sup><br>FW) | Ys<br>(Kg ha<sup>-1</sup>) | STI | YSI | HTI |
| --- | --- | --- | --- | --- | --- | --- | --- | --- | --- | --- | --- |
| F3 |  |  |  |  |  |  |  |  |  |  |  |
|  | Chla | 1.00 | -0.13<sup>ns</sup> | 0.62** | 0.24* | 0.75** | 0.21* | 0.11<sup>ns</sup> | 0.11<sup>ns</sup> | 0.04<sup>ns</sup> | -0.03<sup>ns</sup> |
|  | Chlb |  | 1.00 | 0.69** | -0.69** | -0.11<sup>ns</sup> | 0.16<sup>ns</sup> | 0.24* | 0.26* | 0.17<sup>ns</sup> | 0.11<sup>ns</sup> |
|  | Chla+b |  |  | 1.00 | -0.37** | 0.46** | 0.28* | 0.27* | 0.28* | 0.16<sup>ns</sup> | 0.06<sup>ns</sup> |
|  | Chla/b |  |  |  | 1.00 | 0.05<sup>ns</sup> | -0.04<sup>ns</sup> | -0.11<sup>ns</sup> | -0.12<sup>ns</sup> | -0.10<sup>ns</sup> | -0.05<sup>ns</sup> |
|  | Car |  |  |  |  | 1.00 | 0.01<sup>ns</sup> | -0.15<sup>ns</sup> | -0.15<sup>ns</sup> | -0.10<sup>ns</sup> | -0.11<sup>ns</sup> |
|  | Pro |  |  |  |  |  | 1.00 | 0.43** | 0.41** | 0.38** | 0.18<sup>ns</sup> |
| BC1F2 |  |  |  |  |  |  |  |  |  |  |  |
|  | Chla | 1.00 | -0.19 | 0.60** | 0.35** | 0.67** | 0.17<sup>ns</sup> | 0.04<sup>ns</sup> | -0.01<sup>ns</sup> | 0.30* | 0.31** |
|  | Chlb |  | 1.00 | 0.66** | -0.80** | -0.29* | 0.005<sup>ns</sup> | 0.03<sup>ns</sup> | 0.001<sup>ns</sup> | 0.22<sup>ns</sup> | 0.18<sup>ns</sup> |
|  | Chla+b |  |  | 1.00 | -0.39** | 0.28* | 0.13<sup>ns</sup> | 0.06<sup>ns</sup> | -0.01<sup>ns</sup> | 0.41** | 0.38** |
|  | Chla/b |  |  |  | 1.00 | 0.24* | 0.11<sup>ns</sup> | 0.10<sup>ns</sup> | 0.09<sup>ns</sup> | -0.04<sup>ns</sup> | -0.02<sup>ns</sup> |
|  | Car |  |  |  |  | 1.00 | -0.02<sup>ns</sup> | -0.03<sup>ns</sup> | -0.06<sup>ns</sup> | 0.13<sup>ns</sup> | 0.11<sup>ns</sup> |
|  | Pro |  |  |  |  |  | 1.00 | 0.55** | 0.53** | 0.16<sup>ns</sup> | 0.08<sup>ns</sup> |

Abbreviations: Chla, chlorophyll a content; Chlb, chlorophyll b content; Chla+b, total chlorophyll content; Chla/b, ratio of chlorophyll a to chlorophyll b; Car, carotenoid content; Pro, proline content; Ys: grain yield under high-temperature stress conditions; STI: stress tolerance index; YSI: yield stability index; HTI, heat tolerance index.
†\* and \*\* are significant at 0.05 and 0.01 probability level, respectively. ns, non-significant.

PCA was performed using 14 variables to discriminate the distribution patterns of the families in response to growth conditions and to realize the relationships of families with variables (traits and tolerance indices) under high-temperature stress conditions. The results showed two principal components: Eigenvalues >1 accounting for 36.57% and 19.59% of the variance in the F3 families, respectively (Figure 4). Rely on the multiple traits that correlated strongly with tolerance indices, the superior high-temperature tolerant families should have high PC1 and moderate PC2 in the F3 generation (Figure 4). Accordingly, families no. 8, 9, 10, 20, 21, 60, 62, 83, 84, and 85, could be screened as the most tolerant F3 families to high-temperature stress. In BC1F2 generation, the first component justified 30.87% of the variance, whereas the second component accounted for 22.31% (Figure 5). The PC1 was positively correlated to all traits and tolerance indices except DA in the BC1F2 families. On the other hand, PC2 had a positive association with RGP, NT, total chlorophyll content, and two yield-stability-based indices (YSI and HTI), whereas it showed a negative correlation with NGS, GW, YS, and STI (Figure 5). Thus the BC1F2 families no. 11, 26, 27, 42, and 61 were characterized by the highest yield stability when grown under normal and high-temperature stress conditions (Figure 5). Besides, the families having high PC1 and low PC2 (no. 25, 28, 44, 55, 64, 65, and 66) could be suggested as the most heat-tolerant BC1F2 families.

**FIGURE 4.**
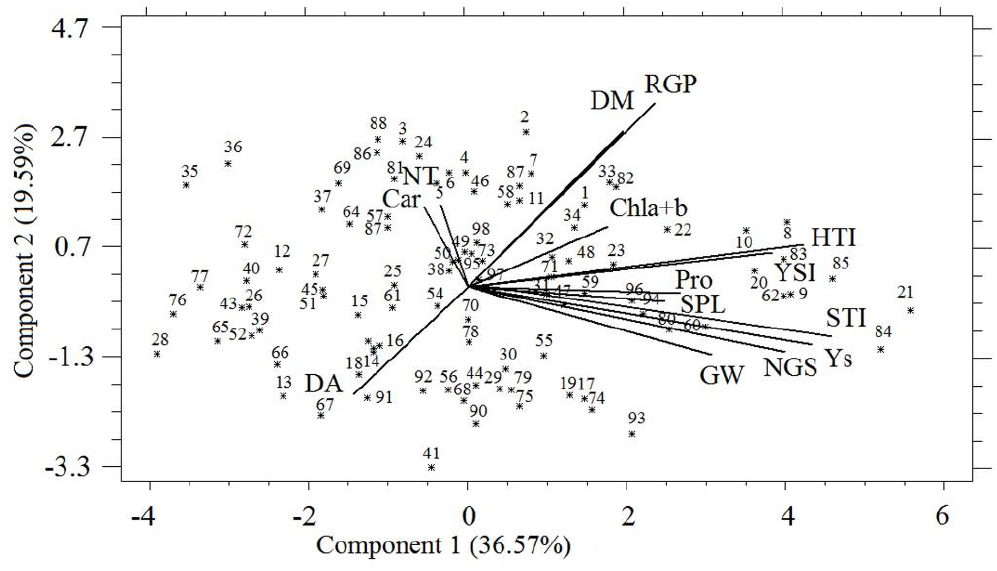
Biplot of PC1 versus PC2 based on the mean values obtained from each trait for 98 F3 barley families. Car, carotenoid content; Chla+b, total chlorophyll content; DA, days to anthesis; DM, days to maturity; GW, 1000-grain weight; HTI: heat tolerance index; NGS, number of grain per spike; NT, number of fertile tillers; Pro, proline content; RGP, reproductive growth period; SPL, spike length; STI, stress tolerance index; Ys, grain yield under high-temperature stress conditions; YSI, yield stability index.

**FIGURE 5.**
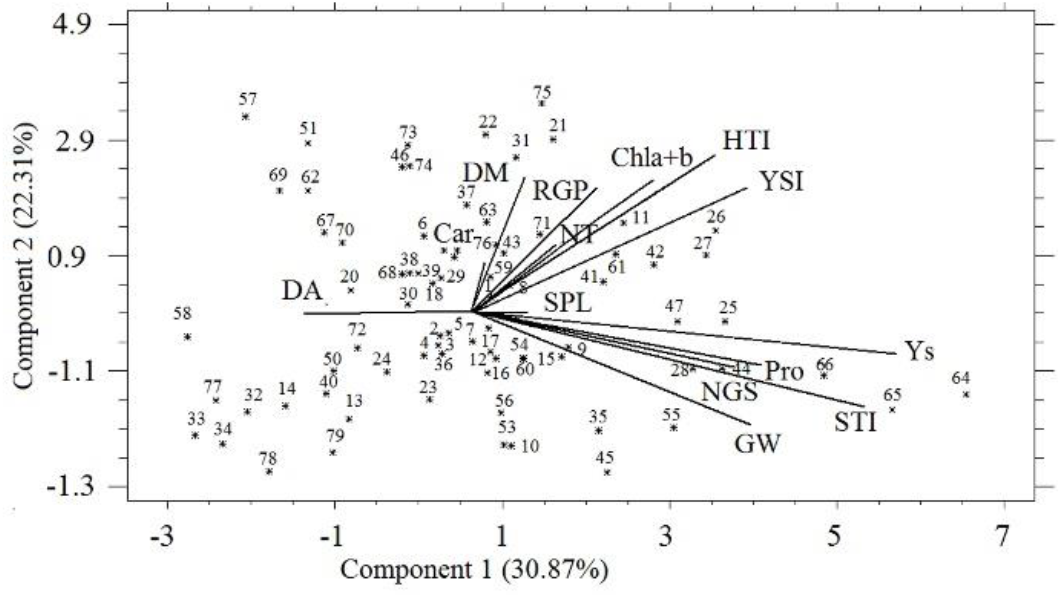
Biplot of PC1 versus PC2 based on the mean values obtained from each trait for 79 BC1F2 barley families. Car, carotenoid content; Chla+b, total chlorophyll content; DA, days to anthesis; DM, days to maturity; GW, 1000-grain weight; HTI: heat tolerance index; NGS, number of grain per spike; NT, number of fertile tillers; Pro, proline content; RGP, reproductive growth period; SPL, spike length; STI, stress tolerance index; Ys, grain yield under high-temperature stress conditions; YSI, yield stability index.

A three-dimensional graph was plotted based on Ys, STI, and YSI to discriminate the most heat-tolerant hybrid families (data not shown). The highest STI, YSI, and grain yield under high-temperature stress conditions were observed in families no.8, 9, 10, 21, 83, 84, 85, and 93 in the F3 generation. Likewise, BC1F2 families no. 25, 27, 44, 55, 64, 65, and 66 had the highest grain yield under high-temperature stress, followed by the higher STI and YSI values.

## 4 | DISCUSSION

Winter cereal crops are generally constrained by high-temperature stress at their terminal growth stage due to the detrimental effects on their reproductive organs and the limited carbon availability during their grain-filling period. It has been postulated compared to their cultivated counterparts, wild cereal species that distributed over a vast area from the Mediterranean regions to central Asia might contain different tolerance genes that evolve in various mechanisms for abiotic stress tolerance (Rampino et al., 2018). This hypothesis is confirmed by our results that wild barley exhibits a rich genetic variation for high-temperature tolerance. Likewise, Hu et al. (2019) reported a high genetic variation for heading dates in wild barley in comparison with cultivated ones in three different sowing dates. It has also been hypothesized that *H. spontaneum* that has been evolved efficient high-temperature tolerance strategies to adaption the hot climate in south-west Iran could be exploited as a rich genetic resource for improving cultivated barley concerning its heat stress tolerance. Recently, this rich genetic resource of *H*. *spontaneum* germplasm originated from West-Iran has been demonstrated to underlie the physiological basis of its tolerance to both heat (Bahrami et al., 2019) and salinity (Ebrahim et al., 2020) stresses.

Late sowing in barley shifted the plant’s reproductive period to times of high-temperature stress. This incidence is in agreement with that reported in wheat (Dwivedi et al., 2017). Reduction in grain yield might be attributed to pollen infertility and seed abortion due to terminal high temperatures during the grain-filling stage (Rezaei et al., 2010; Bita & Gerats, 2013; Jha et al., 2014). Although the *H. spontaneum* germplasm did not exhibit high yield productivity under either treatment, it showed less susceptibility to thermal stress evidenced by its much lower yield loss than those observed in barley cultivars. These findings are consistent with those of Barati et al. (2018), who reported that, compared to the cultivated ones, the *H. spontaneum* genotypes had been less affected by drought stress. Weaker effects of high-temperature stress were also observed in the present study on RGP, NGS, and GW than those on the other traits in the wild genotypes compared with the cultivated ones. It might be, therefore, concluded that the higher reductions in grain yield due to heat stress could have been caused by the shortened reproduction period and the declining yield components. These results, in turn, imply that the heat escape by shortening the life cycle to avoid terminal high temperatures led to reduced yield in both cultivated and wild barley. Our finding of a two-fold reduction in RGP due to heat stress in the cultivated barley implies using ingenuous-focused coping strategies such as escape/avoidance rather than adaptive-focused coping strategy. Lohani et al. (2020) noted that high temperatures during anthesis caused the seed filling duration to shorten, which resulted in smaller grain size and, thereby, a yield decline. Likewise, Zheng et al. (2015) reported the phenological acceleration in wheat under high-temperature stress led to reduced photosynthesis due to smaller leaves and fewer tillers. There is hence a trade-off between the reproduction growth period and tolerance to high-temperature stress. Therefore, uncoupling underlying mechanisms and genetic linkages would enhance breeding new cultivars without the drawback of increased sensitivity.

Since the genes involved in grain yield components are expressed at different developmental stages, they are influenced differently by variations in the environment. In the current study, the lower losses of grain yield in the wild genotypes could not only be attributed to keeping reproduction growth period but also to yield components stability during terminal high temperatures. Jedmowski et al. (2015) found that heat stress had no significant effect on the total number of spikes per plant in *H. spontaneum* but that the number of seeds per spike and seed weight declined under high-temperatures stress conditions. In the wild genotypes, fertile tiller production and percentage of fertile florets exhibited more substantial effect of terminal high temperatures than did seed size. These results are consistent with those reported by Jedmowski and coworkers (2015), who noted that disturbances in floral development due to heat stress affected grain productivity in wild barley.

Understanding tolerance responses of plants to high temperatures is imperative for maintaining yield in the context of plasticity an adaptation to seasonally fluctuating environments in the face of climate change. (Given the fact that successful breeding programs can be measured against proper indices, an ideal tolerance index should have a high discriminative power to identify superior genotypes with long-term yield stability (Bahrami et al., 2014). The need to screen for stress-tolerant germplasm has, therefore, urged plant breeders to search for reliable indices. A modified tolerance index was proposed in this paper to incorporate the most critical indicators of performance, including yield stability and yield loss to stress. The results showed a high variation in the wild barley germplasm, and a discriminative screening judgment was made using each of the five indices.

Despite the much higher yield of the cultivated genotypes compared to the wild ones studied, the highest yield reduction due to thermal stress, or the highest TOL, was found to belong to four barley cultivars. In addition, the highest STI values were recorded for the three cultivated barley genotypes of ‘Reyhan’, ‘Nosrat’, and ‘Fajr30’. The results followed an expected pattern with an STI index that took into account the yields in both environmental treatments. These findings are consistent with those of a recent study showing higher TOL and STI values in cultivated genotypes (*H. vulgare)* compared to wild ones (Barati et al., 2019) subjected to drought stress. Indeed, each of the tolerance indices used imposes important restrictions on the screening process so that the strategies used may not be uniformly practicable. It is, therefore, essential to employ a combination of multiple supplementary indices that will not overemphasize only certain aspects of the genotypic attributes at the cost of a reasonable overall evaluation of genotype performance. In this regard, the strong correlations among Ys, STI, YSI, and HTI were used to identify the wild genotypes no.1, 2, 15, 17, 21, and 41 with the highest STI and YSI, and HTI values as the ones tolerant to high-temperature stress. On the other hand, genotypes no.6, 25, 35, and 43 with the highest SSI values were identified as the most susceptible to thermal stress among wild genotypes.

PCA is a robust statistical procedure to reduce the dimensions of the variables and to divulge constructive evidence-driven feedback from a highly-correlated dataset (Banerjee et al., 2020). The two-dimensional plot obtained from the two PC scores proved YSI and HTI as the most potent yield-stability based indices to identify tolerant wild genotypes capable of maximum sustainable yield. Under high-temperature stress, the positive correlations among DM, RGP, SPL, and NT concentrated in the optimum region in the plot; i.e., they had high PC1 and PC2 scores, suggesting that these traits were important in defining high-temperature tolerant wild genotypes. Using a combination of drought tolerance indices, Bahrami et al. (2014) and Barati et al. (2019) identified drought-tolerant genotypes in safflower and wild barley, respectively. Our results showed that the high-temperature tolerant wild barley genotypes exhibited taller spike heights, more grain per spike, and sufficient grain-filling duration when exposed to high temperatures in the reproductive stage. Thus as the grain (number and weight per spike) influences significantly from grain-filling duration and contributes significantly to final yield, reducing its number to enable heat tolerance or escape is likely to result in a substantial trade-off with yield. Chaudhary et al. (2020) noted that grain-filling duration is a vulnerable stage to high-temperature which directly associated with grain number and yield. These findings are consistent with those reported by Bergkamp et al. (2018) on wheat, indicating that terminal heat stress can shorten the grain-filling period via accelerated grain-filling rate, thereby causing a limited amount of carbohydrates transferred from the source tissues to the grains and reducing grain yield.

Moreover, the results indicated that shortened life cycle to escape terminal high temperatures is a common defense mechanism adopted by both cultivated and wild genotypes of barley. This is particularly important given the negative effect of drastically grain yield performance but the positive one of dramatically enhanced adaptation to environment. In the present study, it was found that YSI and HTI could be exploited to identify the cultivated and wild barley genotypes with the highest thermal tolerance by making a reasonable compromise between yield and its stability. It was also found that the overall performance of wild populations could be viewed as an evolutionary compromise between two advantageous features of plant evolution - that is, increased productivity in an extremely stressful ecosystem and its long-term sustainability.

Barley wild relatives could be a source of desirable traits associated with abiotic stress tolerance, which might be exploited in conventional breeding programs (Bahrami et al., 2019; Baratiet al., 2019; Ebrahim et al., 2020). Using hybridization between wild and domesticated barley provides more genetic variation to improve abiotic stress adaptability in cultivated genotypes (El-Hashash & El-Absy, 2019). In the current study, the wide range of variation observed among F3 and BC1F2 families for agronomic and physiological traits. This in turn illustrated how introgression of wild parent genome into cultivated barley is driving more diversity in the traits contributing to high-temperature tolerance. Moreover, results also showed that some progenies exceeded well beyond that of the parents for grain yield in F3 and BC1F2 families exhibiting transgressive segregation.

In the current study, BC1F2 families exhibited a higher grain weight and yield, followed by the lowest reduction in NT and GW compared to the F3 generation and even both parents under high-temperature stress. Furthermore, BC1F2 families showed higher stability than cultivated parent under high-temperature. Although more backcross should be carried out to remove undesirable wild attributes, the present study has demonstrated that even with one backcross only, the stability of progenies is improved efficiently.

Accordingly, PCA results were in agreement with those observed in wheat (Abou-Elwafa & Shehzad, 2021) and barley (Barati et al., 2019), indicating that yield-based tolerance indices can help breeders in choosing cultivated and wild genotypes as well as their derived families for abiotic stress tolerance, either when used alone or in combination with agronomic and physiological parameters. In the current study, NGS and GW were the most effective yield components for screening superior high-temperature tolerant F3 and BC1F2 families. HTI and YSI were more successful in discriminating the families with higher yield stability under high-temperature stress conditions. Therefore, these findings support the idea of a combined implication of tolerance indices and yield components for the selection of genetic materials with sustainable production. Overall, the comparisons of F3 and BC1F2 families with their cultivated parent indicated that the high-temperature tolerance from the wild parent has been partially inherited to the descendants.

## 5 | CONCLUSION

A high level of variability exists in the Iranian *H. spontaneum* germplasm for agronomic traits leading to better yield stability due to tolerance to thermal stress. The introgressed lines also showed the positive contributions of this variability. The ingenuous-focused strategies like escape/avoidance are being used primarily to cope with heat stress by cultivated barley, while adaptive-focused coping strategies such as tolerance are being implemented by wild barley. Results also indicated that indirect selection for grain yield using a combination of tolerance indices and agronomic attributes could be effective to select in the high-temperature tolerant lines of barley. Ultimately, a single yield-based index cannot be employed to allow identification of the sustainable high-yielding lines for growth in both normal and stress conditions because it cannot afford the best opportunity of combining high yield with yield stability. Alternatively, the use of more than one tolerance index or combined use of tolerance index and agronomic/physiological attributes should reduce these obstacles and facilitate accurate and reproducible identification of the most stable and high yielding genotypes.

## ACKNOWLEDGEMENT

This work was supported by funds from the Iran National Science Foundation (INSF) under Grant No. 96002328.

## AUTHOR CONTRIBUTIONS

AA collected the seeds and contributed significantly to the study design. This study was conducted and analyzed by FB with supervision from AA and MR. FB wrote the manuscript with significant input from AA and MR.

## Abbreviations

Car: carotenoid content
Chla: chlorophyll a content
Chlb: chlorophyll b content
Chla+b: total chlorophyll content
Chla/b: ratio of chlorophyll a to chlorophyll b
DA: days to anthesis
DM: days to maturity
GW: 1000-grain weight
HTI: heat tolerance index
NGS: number of grain per spike
NT: number of fertile tillers
PH: plant height
Pro: proline content
RGP: reproductive growth period
SPL: spike length
SSI: stress susceptibility index
STI: stress tolerance index
TOL: tolerance index
Yp: grain yield under normal conditions
Ys: grain yield under high-temperature stress conditions
YSI: yield stability index.

